# Hierarchical Discovery of Large-scale and Focal Copy Number Alterations in Low-coverage Cancer Genomes

**DOI:** 10.1101/639294

**Authors:** Ahmed Ibrahim Samir Khalil, Costerwell Khyriem, Anupam Chattopadhyay, Amartya Sanyal

## Abstract

**Motivation:** Detection of copy number alterations (CNA) is critical to understand genetic diversity, genome evolution and pathological conditions such as cancer. Cancer genomes are plagued with widespread multi-level structural aberrations of chromosomes that pose challenges to discover CNAs of different length scales with distinct biological origin and function. Although several tools are available to identify CNAs using read depth (RD) of coverage, they fail to distinguish between large-scale and focal alterations due to inaccurate modeling of the RD signal of cancer genomes. These tools are also affected by RD signal variations, pronounced in low-coverage data, which significantly inflate false detection of change points and inaccurate CNA calling.

**Results:** We have developed CNAtra to hierarchically discover and classify ‘large-scale’ and ‘focal’ copy number gain/loss from whole-genome sequencing (WGS) data. CNAtra provides an analytical and visualization framework for CNV profiling using single sequencing sample. CNAtra first utilizes multimodal distribution to estimate the copy number (CN) reference from the complex RD profile of the cancer genome. We utilized Savitzy-Golay filter and Modified Varri segmentation to capture the change points. We then developed a CN state-driven merging algorithm to identify the large segments with distinct copy number. Next, focal alterations were identified in each large segment using coverage-based thresholding to mitigate the adverse effects of signal variations. We tested CNAtra calls using experimentally verified segmental aneuploidies and focal alterations which confirmed CNAtra’s ability to detect and distinguish the two alteration phenomena. We used realistic simulated data for benchmarking the performance of CNAtra against other detection tools where we artificially spiked-in CNAs in the original cancer profiles. We found that CNAtra is superior in terms of precision, recall, and *f-measure*. CNAtra shows the highest sensitivity of 93% and 97% for detecting focal and large-scale alterations respectively. Visual inspection of CNAs showed that CNAtra is the most robust detection tool for low-coverage cancer data.

**Availability and implementation:** CNAtra is an open source software implemented in MATLAB, and is available at https://github.com/AISKhalil/CNAtra

## 1. INTRODUCTION

Copy number variation (CNV) is an essential genetic variation that leads to the change in the number of copies of genomic regions in comparison to the reference genome. CNVs include small-scale (<100 bp) insertions or deletions (indels) and copy number alterations (CNAs). CNA events are ‘relative’ gain or loss of DNA compared to the reference sample(s) or assembly that are between 1 kb and 3 Mb in size [1]. In cancer, the overwhelming extent of CNA size distribution resulted in their further classification into large-scale (>25% of chromosome arm) or focal events [2, 3]. First, *microscopic level* large-scale copy number variations (LCVs) concern chromosomal abnormalities that can be cytogenetically detected such as segmental aneuploidy. LCVs may also represent polymorphic variations among individuals [4]. Second, the *submicroscopic level* ‘focal’ chromosomal aberrations can range between several kb to a few Mb in size containing a small number of genes [5] believed to harbor important oncotargets. Therefore, CNV detection methodology needs to be tuned to identify both large-scale and focal events and should include procedures to distinguish them. Both LCVs and focal alterations (FAs) occurring at different genomic length scale and amplitude, hold tremendous value for cancer diagnostics and therapeutics. Naturally, their accurate detection is crucial for gaining insights on their origin and biological context.

Numerous computational tools have been developed for CNV detection using next-generation sequencing (NGS) data [6–8]. These tools are in strong demand as they are central to CNV prediction from personalized genomes or individual tumor biopsy for clinical decision-making. Most common NGS-based detection tools use depth of coverage from WGS data to identify CNAs by modeling the read depth (RD) signal either from an individual sample or taking advantage of matched normal or reference samples [6]. In practical scenarios, matched normal samples from same tissue are difficult to procure. Therefore, single-sample tools that do not require matched controls are more applicable in clinical settings and in case of cell lines. Several tools have been presented over the years that utilize single sequencing sample for CNV detection [9–16]. They are built on different assumptions of the underlying probabilistic distribution and percentage of variations. However, in contrast to normal genomes, cancer cells are plagued with large-scale segmental aneuploidies which may lead to inaccurate estimation of the CN reference. Moreover, disregarding the distinction of large-scale and focal events, these CNV calling approaches suffer from oversegmentation of LCVs and erroneous calling of FAs. Therefore, none of the available tools can capture the entire spectrum of copy number aberrations in cancer cells and distinguish between the LCVs and FAs. Moreover, RD based CNV detection tools are adversely affected by low-coverage data resulted in overdispersion and short-term variations such as wave artifacts [17–19]. In such scenarios, statistics-based segmentation [9, 14, 20] and CNV calling lead to either in false segmentation or missing the focal alterations. However, low-coverage NGS data is still more efficient than array-based data for CNV analysis [21].

To address these challenges, we developed CNAtra (**<u>C</u>**opy **<u>N</u>**umber **<u>A</u>**lteration (detection) **<u>t</u>**hrough **<u>r</u>**ead depth **<u>a</u>**nalysis), a MATLAB-based hierarchical computational framework for the sensitive and robust detection of LCVs and FAs. CNAtra is built on the fundamental assumption that most genomic regions of any cell are centralized on copy number state of positive integer values. CNAtra empirically models the RD signal based on a multimodal distribution and estimate the CN reference. This approach allows us to define the accurate ‘interval’ of copy number states which aids in identifying segmental aneuploidies (and FAs within them) in a robust manner largely unaffected by coverage, percentage variation, and wave artifacts. For this, we first applied a robust signal-processing technique of univariate time series to identify significant change points of the RD signal. These change points are used for assembling large segments based on CN state and for identifying the candidate focal alterations. In addition, for handling the overdispersion problem of low-coverage data, we incorporated coverage-based thresholding parameters besides the conventional statistical test to identify the significant FAs. CNAtra also provides an interactive platform to visualize and manually inspect the complete CNA profile and accessory information for further validation, interpretation and application of CNA calls. We successfully verified CNAtra results using experimentally validated segmental aneuploidies and focal amplifications/deletions across several cancer cells. We then benchmarked the performance of CNAtra against five single-sample CNV detection tools using realistic simulated data. For this, we randomly spiked-in large and focal copy number gain/loss in original cancer genomes. The results showed the ability of CNAtra to resolve complex CNV profiles into LCVs and FAs with the highest *f-measure*. Manual review and visualization also verified the advantage of CNAtra over other tools.

## 2. METHODS

The detailed description of methods is provided as Extended Methods under Supplementary Information. Here, we briefly explain the CNAtra framework, coverage-based parameter calibration, and generation of simulated data.

### 2.1 CNAtra framework

CNAtra has been developed to detect LCVs and FAs in cancer genomes hierarchically. CNAtra is a MATLAB-based single-sample CNA discovery tool particularly adapted for low-coverage cancer genomes. CNAtra comprises two modules – RD calculator and CNV caller. In the RD calculator, we compute the RD signal as base count frequency at 1 kb bin from the input WGS data after initial read filtering steps. One kb bin allows us to fine-tune the tool to precisely define the boundaries for both LCVs and FAs. We then correct the RD signal for systematic biases due to GC content (isochore normalization) and low-mappability regions. The CNV caller module constitutes the hierarchical framework to delineate the multi-level alterations in the cancer genomes. We first compute the CN reference by fitting a multimodal distribution over the RD frequency histogram. Second, we use a multi-step framework to identify ‘large’ segments with distinct CN state. Third, we discover candidate peaks of ‘focal’ amplification and deletions in each CN-defined large segment.

#### 2.1.1 Estimation of copy number reference

We utilize a multimodal distribution for computing CN reference (2N) and higher CN states (3N, 4N, …) based on the ploidy level of the given cell line. Our multimodal distribution is defined as the summation of normal distributions of different probabilities centered at different CN states. CN reference is computed as the RD signal per bin that achieves the best correlation between the copy number states and the band of equimolar reads observed in the frequency distribution. Whole genome ploidy level (diploid/triploid/tetraploid) can be selected based on prior knowledge. Otherwise, CNAtra can compute the CN reference independently based on a free model.

#### 2.1.2 Detection of iso-copy number block (IB)

We apply the Savitzky–Golay smoothing filter [22] to eliminate short-term variations and wave artifacts without affecting the ‘sharp’ signal change points. This filter is not influenced by shifting effect which is important for detecting the true boundaries of segments [23]. We then adapted the Modified Varri method [24] to detect the amplitude-shift points of the RD signal that defines the boundary of primary segments. A subsequent CN state-based merging process combines adjacent initial segments into large contiguous segments. We termed these merged contiguous segments with distinct CN state as **iso-copy number blocks (IBs)**. IBs with CN state different from CN reference are considered as segmental aneuploidy or LCVs.

#### 2.1.3 Identification of focal alteration

Each IB is used as a population of bins for the discovery of FAs. Assuming a normal distribution, we perform the t-test to identify the *statistically significant* FA(s) in each IB. Additionally, we define *high confidence* focal amplifications and deletions using coverage-based thresholding. These thresholds represent the minimum amplitude-shift between the estimated CN of candidate regions and their parent IBs to call FAs. We also filter out the FAs that are in problematic/challenging regions of the genome or if they are smaller than CNAtra resolution. This resolution is the minimum width of FA that can be detected with FDR < 0.05 based on the genome coverage.

### 2.2 Estimation of CNAtra calibration parameters

We employ the exponential regression function to model the relationship between the data coverage and CNAtra parameters such as resolution, amplification, and deletion thresholds. We used datasets of high genome coverage (10x-14x) which include 1000 Genomes Project datasets (HG00119, HG01879, HG00096) and A427 cell line. Then, we computed the optimum values of the analysis parameters of the original and subsampled data. Subsampling was performed using Picard (http://broadinstitute.github.io/picard/) and SAMtools [25]. These values were used for fitting the exponential regression models.

### 2.3 Simulated CNV profile by randomly introducing CNA regions in the real cancer data

We generated simulated CNV profile from real cancer data. Starting from NGS reads of cancer data, we artificially introduced both LCVs and FAs of random copy number and width into cherry-picked chromosomes devoid of any observable large structural variations. For each spiked alteration region, we added or removed the number of reads in proportion to the desired copy number by utilizing actual reads of the targeted region. These spiked-in CNAs provided us with *de facto* ground truth for performance evaluation of CNV calls in terms of CN estimation, accuracy, and precision. We used these simulated data for benchmarking CNAtra against five other CNV detection tools.

### 2.4 Data availability

All the datasets used in this study are publicly available (Supplementary Table 1). We used CNA profiles of cancer cell lines from CCLE [26] and COSMIC [27] databases for performance evaluation of CNAtra.

## 3. RESULTS

### 3.1 Cancer genomes harbor LCV and FA with distinct biological origins

Cancer cells are afflicted with widespread numerical and structural variations of chromosomes [28, 29] which positively correlate with tumor aggressiveness. Cancer cells contain both LCVs and focal alterations having different mechanisms of origin and functional roles. LCVs are whole chromosome or large genomic ‘blocks’ (> 1Mb) with distinct CN states that result from chromosomal instability which results in acquisition of complex genetic makeup by cancer cells [30]. In contrast, focal amplification and deletions emerge as a consequence of adaptive selection events that facilitate selective growth advantage and evolution of malignant cells during tumor development and drug resistance [3]. Focal amplifications generally have high-level gains of oncogenes or anti-apoptotic genes while focal deletions usually involve tumor suppressor or pro-apoptotic genes [31, 32]. Therefore, identification and characterization of these two phenomena can provide vital clues to identify the genomic regions and driver genes involved in carcinogenesis and their roles in cancer evolution. An illustrative example using a coverage plot from WGS data of A427 cell line is provided in Figure 1. LCVs are pervasive genome-wide in A427 which result in the multimodal frequency distribution of RD signal (Fig. 1a). A closer look at chromosome 2 (Fig. 1b) shows that some focal events are interspersed within LCVs creating a complex relationship between them. For example, focal amplifications containing *USP34* and *CCNT2* genes are part of different LCVs in 2p and 2q region respectively (Fig. 1b). Therefore, there can be a complex scenario where a genomic segment may have a hemizygous segmental deletion (LCV) which in turn can contain a focal amplification. In the coverage plot, these LCVs appear as ‘large’ segments, and they are strongly affected by ‘wave artifacts’ (indicated as a blue curve in Fig. 1b,c). Wave artifacts are systematic biases due to deviation from equimolar coverage [33]. On the other hand, focal amplifications and deletions are represented as ‘sharp’ peaks and troughs respectively (Fig. 1b,c). Moreover, RD signals from WGS are also prone to inherent biases associated with NGS owing to genome GC content, low-mappability regions, and coverage-influenced overdispersion of the RD signal. All these biases ultimately complicate the multi-level detection of segmental aneuploidies and FAs. Taken together, it can be concluded that cancer genomes have multi-level CNAs and their RD signal are inherently complex as evidenced by nature and distribution as well as associated systematic biases. Therefore, CNA detection in cancer genomes necessitates the biological understanding of this phenomenon and based on which a step-by-step approach needs to be implemented to delineate the multi-level aberrations one at a time. None of the currently available tools have adequately addressed these multi-level issues *in toto*.

**Fig. 1.**
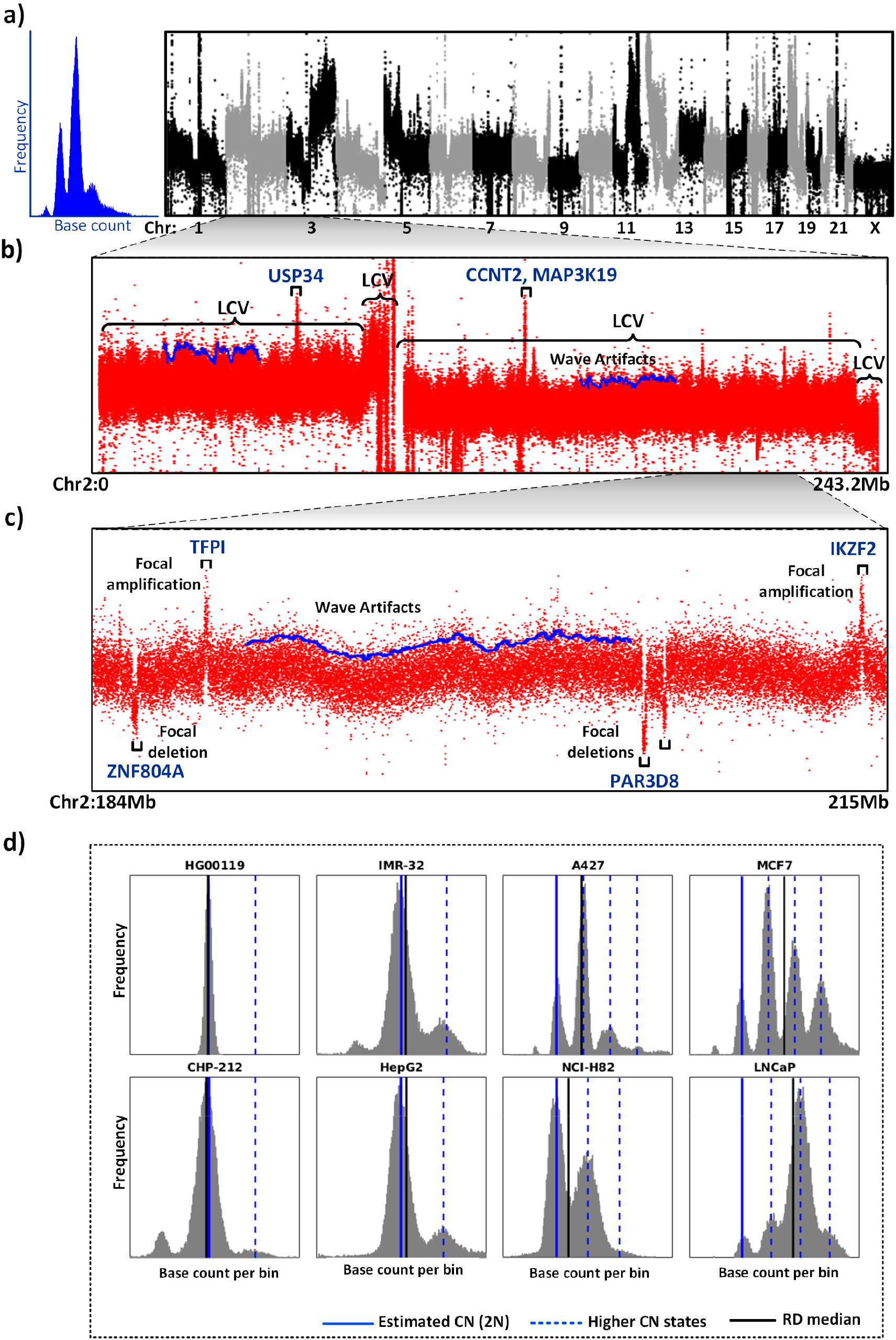
Characterization of RD signal of cancer genomes. **(a)** The RD frequency distribution (left) and genome-wide coverage plot of all chromosomes (right) of A427 cell showing the presence of widespread large-scale segmental aneuploidies. **(b)** Coverage plot of Chr 2 showing LCVs of different copy number states. **(c)** Zoomed in view of Chr 2 (184-215 Mb) shows the presence of focal alterations (both amplification and deletions) inside the LCV. The blue curved line in **(b)** and **(c)** denotes the upper envelope of RD signal showing regions affected by wave artifacts. Few focal amplifications and deletions are indicated. **(d)** Estimation of CN reference (2N) using CNAtra (blue line) and the global median (black line) for normal genome (HG00119) and different cancer cell lines. The dotted blue line denotes the higher CN states (3N, 4N, 5N) based on CNAtra CN reference.

### 3.2 Accurate estimation of CN reference is essential for CNA calling in cancer genomes

In addition to the presence of LCVs, chromsome segregation errors lead to elevated ploidy (at genome or chromosome levels) and karyotype alterations of cancer cells [34]. Human cancers frequently have hyperdiploid, near-triploid or higher ploidy levels [35–37]. All these anomalies manifest as multimodal distribution of the RD signal. We analyzed several cancer cell lines of different ploidy and complexity using publicly available data [35, 38–44] (Supplementary Table 1). Cancer genomes exhibited a complex *multimodal* distribution as opposed to normal genomes (1000 genomes project) which follow a unimodal distribution (Fig. 1d; Supplementary Fig. S1). We also found that medians of RD signal across chromosomes are highly inconsistent for cancer cells (Supplementary Fig. S1). Most single-sample tools assume unimodal distribution of the RD signal and use the global median as CN reference (2N). In contrast, CNAtra utilized a multimodal distribution as a summation of normal distribution of different probability centralized at CN states under a given ploidy assumption (Fig. 2a). This allows accurate estimation of the CN reference. As shown in Figure 1d, our estimated CN reference (2N; solid blue line) and higher CN states (dashed blue line) are coinciding with peaks of the RD signal. However, the median-based CN reference (solid black line) deviates from the ‘actual’ CN reference (solid blue line) by 2.5-87% (Supplementary Table 2) depending on the percentage of LCVs in the cancer cells as opposed to 0.34-0.56% for normal genomes.

**Fig. 2.**
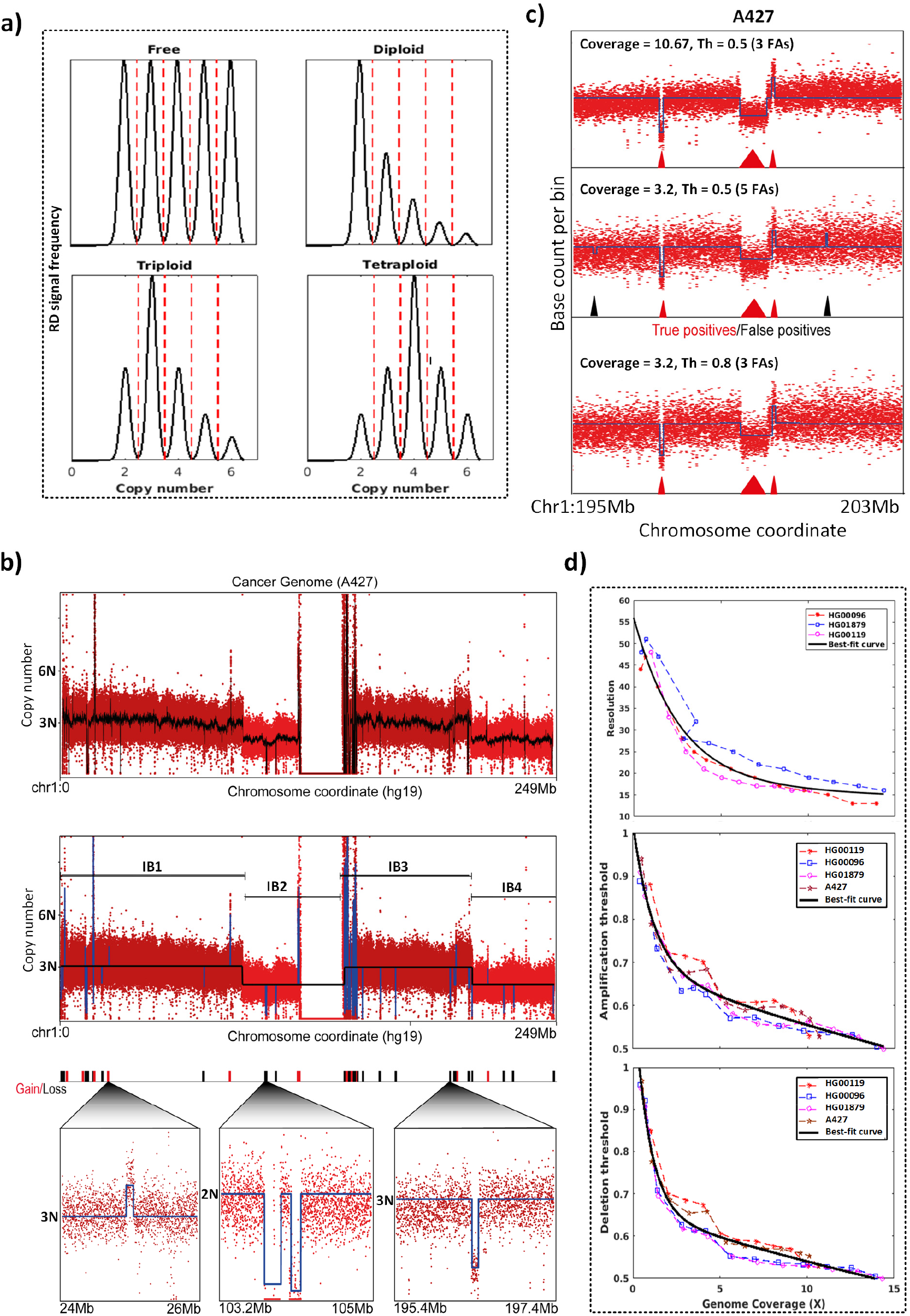
CNAtra solution for multi-level CNA detection in low-coverage data. **(a)** Schematic representation of multimodal distribution under different ploidy assumptions. Dotted red lines denote the boundaries of CN states. **(b)** A hierarchical framework for detecting both LCVs and FAs. The CNAtra approach includes smoothing of RD signal using Savitzky-Golay filter (black line) (top panel), followed by detection of IBs with distinct CN states (middle panel) and then identification of FAs inside each IB (bottom panel). **(c)** Effects of tuning the amplification/deletion thresholds (Th) on the detection of false positive CNAs in a Chr 1 region (hg19 chr1:195-203 Mb) using subsampled data of A427. The red and black triangles represent the true positives and false positives respectively. **(d)** Calibration of CNAtra parameters such as resolution (top panel), amplification threshold (middle panel) and deletion threshold (bottom panel) using high-coverage datasets.

Nevertheless, karyotype or ploidy levels information is not always available. In that case, our free model can still predict the CN reference with a maximum error of 0.44% compared to the actual CN reference given the ploidy information. Therefore, our multimodal approach can be applied universally regardless of the knowledge of the underlying ploidy level or karyotype. Defining the CN reference forms the basis to define the segmental aneuploidies and estimate the copy number states accurately.

### 3.3 Hierarchical framework enables CNAtra to detect and distinguish between LCV and FA

We have taken a pragmatic approach for solving the two major problems associated with cancer genomes- 1) presence of LCV and 2) systematic biases such as overdispersion and wave artifacts which are pronounced in low-coverage data. Savitzky–Golay smoothing filter successfully attenuated the wave artifacts and signal variability to identify the primary segments (Fig. 2b top panel black line). However, CNV detection methods based solely on segmentation may suffer from false segmentation or oversegmentation. For example, neighboring segments may represent CNVs belonging to same CN state which has no biological basis (false segmentation). In addition, a LCV region can be falsely ‘oversegmented’ into several regions with the same copy number due to the presence of focal amplifications and deletions. This may lead to failure in capturing the entire LCV as a single event. We solved this by the IB assembly algorithm which successfully merges primary segments with the same copy number to define the IBs (Fig. 2b middle panel). IBs represent distinct (unimodal) peaks in the multimodal RD signal distribution centered on its calculated CN state. IBs with CN state different from CN reference are considered as segmental aneuploidy or LCVs.

After defining the IBs, we found that RD signal from each IB follows the normal distribution using Q-Q plot and Kolmogorov–Smirnov test with a reasonably good approximation (5% significance level) (Supplementary Fig. S2). Therefore, we use each IB as a population of bins for the discovery of *statistically significant* FAs **(Class 1)** using the t-test. However, due to overdispersion of RD signal in cancer genomes, statistical tests may reject the null hypothesis of long segments with a small mean difference (particularly in the presence of wave artifacts) resulting in many false positives. Therefore, we employed coverage-based thresholding to define *high confidence* **(Class 2)** FAs based on the local CN reference of the parent IB (Fig. 2b bottom panel).

### 3.4 Coverage-based thresholding enables the detection of high confidence CNAs in low-coverage datasets

Dispersion of RD signal is inversely proportional to the depth of coverage. Therefore, CNV detection tools usually recommend using large bins (coverage-driven bin-size approach) to avoid false detection of CNAs in low-coverage datasets. However, this approach may result in missing significant focal alterations. In contrast, we propose to adapt the coverage-driven thresholding approach instead of increasing the bin-size. This allows the identification of focal CNAs without increasing the false positives.

Theoretically, any ‘candidate’ focal alteration with amplitude-shift > 0.5N (threshold) from the CN of its parent IB can be identified as significant focal alteration since it belongs to another copy number state (Supplementary Fig. S3). However, low-coverage data suffer from higher variability resulting in false-calling of focal CNAs with the same threshold (Fig. 2c). Therefore, we define the coverage-based parameters (resolution, amplification and deletion thresholds) to adapt with this overdispersion in low-coverage data (Fig. 2d). CNAtra is calibrated using several WGS datasets of high coverage. We utilized negative exponential regression for modeling the relation between sequencing coverage and the coverage-based parameters. Our thresholding parameter enables a user to strike a balance between false positives and false negatives. For example, using the same threshold of 0.5N, the subsampling of A427 data to 3.2x coverage yields more false positives compared to the original 10.67x coverage (Fig. 2c top and middle panel). Increasing the threshold gets rid of these false positives (Fig. 2c bottom panel). Therefore, the advantage of coverage-based parameter tuning makes the CNAtra results more robust at different data coverage.

### 3.5 CNAtra detects experimentally validated LCVs and FAs across cell lines

We evaluated the ability of CNAtra to detect and distinguish both large-scale and focal events using validated data as ‘ground truth’ from multiple cancer datasets. For LCVs, CNAtra confirms the complete genetic profile of LCVs of HepG2 reported earlier using array CGH (comparative genomic hybridization) analysis [41]. CNAtra correctly identified the whole chromosome (2, 16 and 20) and segmental (involving Chr 6, 14, 17) gains (Fig. 3a, Supplementary Fig. S4). Moreover, CNAtra precisely identified the well-known mono-allelic 1p deletion in neuroblastoma cell lines (IMR-32 and CHP-212) [45] (Fig. 3b). In both these cells, CNAtra called 1p deleted region as a single LCV event with correct CN estimation (CN=1). On the other hand, CNAtra also successfully detected the *well-known* focal amplifications of *MYC* (NCI-H82) [46] and *MYCN* (CHP-212, IMR-32) [47] loci as well as homozygous focal deletions of *BANK1/4q24* (NCI-H82) [48], *LKB1/STK11* (A427) [49], and *p16INK4a/CDKN2A* (A427) [50] loci (Fig. 3c; Supplementary Fig. S4).

**Fig. 3.**
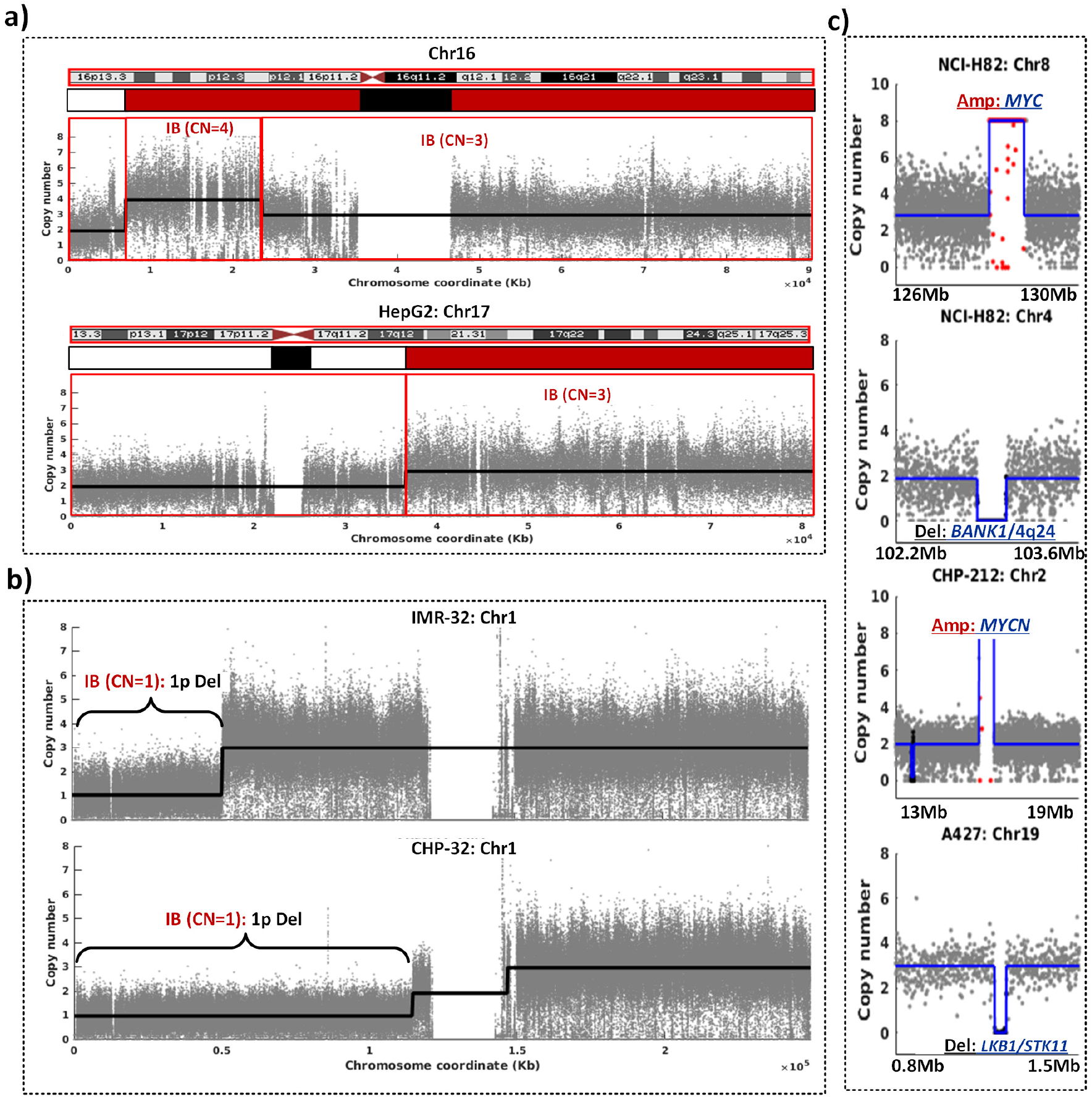
CNAtra validation of well-known LCVs and FAs. **(a)** CNAtra correctly detects the large segmental aneuploidies of Chr 16 (top) and Chr 17 (bottom) of HepG2 cell line described earlier using array CGH (shown as red bars below the idiograms). The black bar denotes the centromere region while the white bar represents normal region (2N). **(b)** CNAtra identifies monoallelic 1p deletion in both IMR-32 and CHP-212 neuroblastoma cells. **(c)** CNAtra detects focal amplifications (*MYC, MYCN*) and focal deletions (*BANK1*/4q24 and *LKB1/STK11*) in respective cancer cell lines.

### 3.6 CNAtra confirms the CNV profiles identified by SNP array in CCLE and COSMIC database using low-coverage NGS data

In order to estimate our performance in a global manner, we used genome amplification data of CHP-212 and NCI-H82 cells which are available from both CCLE [26] and COSMIC [27] databases from SNP array data (Supplementary Table 3). In the COSMIC database, CNAs were called using PICNIC software [51] while CCLE calls were based on circular binary segmentation (CBS) approach [20]. We applied CNAtra on the low-coverage NGS data of CHP212 (1.4x) and NCI-H82 (0.31x) and compared our calls. In COSMIC data, the CNAs have been called in a gene-centered manner where each region is associated with one or more genes. COSMIC data has 6 and 13 amplified regions for the CHP-212 and NCI-H82 cells respectively. We found that out of 6 amplified regions in CHP-212, CNAtra identified two as focal CNVs and other four regions inside a single LCV (Supplementary Fig. S5a). Similarly, for NCI-H82, 8 out of 13 CNVs have been detected as focal events, and three are embedded in two LCVs. In comparison, we found that CCLE detected 11 and 99 amplified regions in CHP-212 and NCI-H82 cells respectively. In CHP-212, CNAtra detected 10 out of 11 amplified regions as four focal events, three LCVs and, the rest three are embedded in one LCV. Similarly, 78 out of 99 amplified regions were detected by CNAtra for NCI-H82. Out of 78, CNAtra identified 11 as FAs, 7 LCVs and rest 60 are part of 19 LCVs. This suggests that CCLE calls are segmenting LCVs into smaller CNA segments (Supplementary Fig. S5b). Taken together, CNAtra detected 89.5% COSMIC and 80% of CCLE calls.

Surprisingly, there are only two CHP-212 and four NCI-H82 calls that were detected by both COSMIC and CCLE as consensus amplification regions suggesting poor concordance to identify CNV events. These six consensus regions are detected by CNAtra. However, the number of consensus regions is extremely low for a robust assessment of performance. Also, COSMIC and CCLE CNVs do not distinguish between LCVs and FAs. Therefore, the absence of a detailed cancer CNV profile necessitates the usage of realistic simulated data where CNAs can be artificially introduced to use as ground truths for performance evaluation.

### 3.7 CNAtra is the superior tool for detecting focal and large CNAs in low-coverage data using realistic simulated data

In the absence of extensively validated datasets of both LCVs and FAs, we relied on the simulated CNV profile. However, a simple simulation may not capture the inherent biases of the RD signal of cancer genomes. Therefore, we developed a novel approach to manipulate the original WGS reads of cancer genome to randomly introduce FAs embedded within the LCVs maintaining the inherent CNA features and complexities of the RD signal (Fig. 4a).

**Fig. 4.**
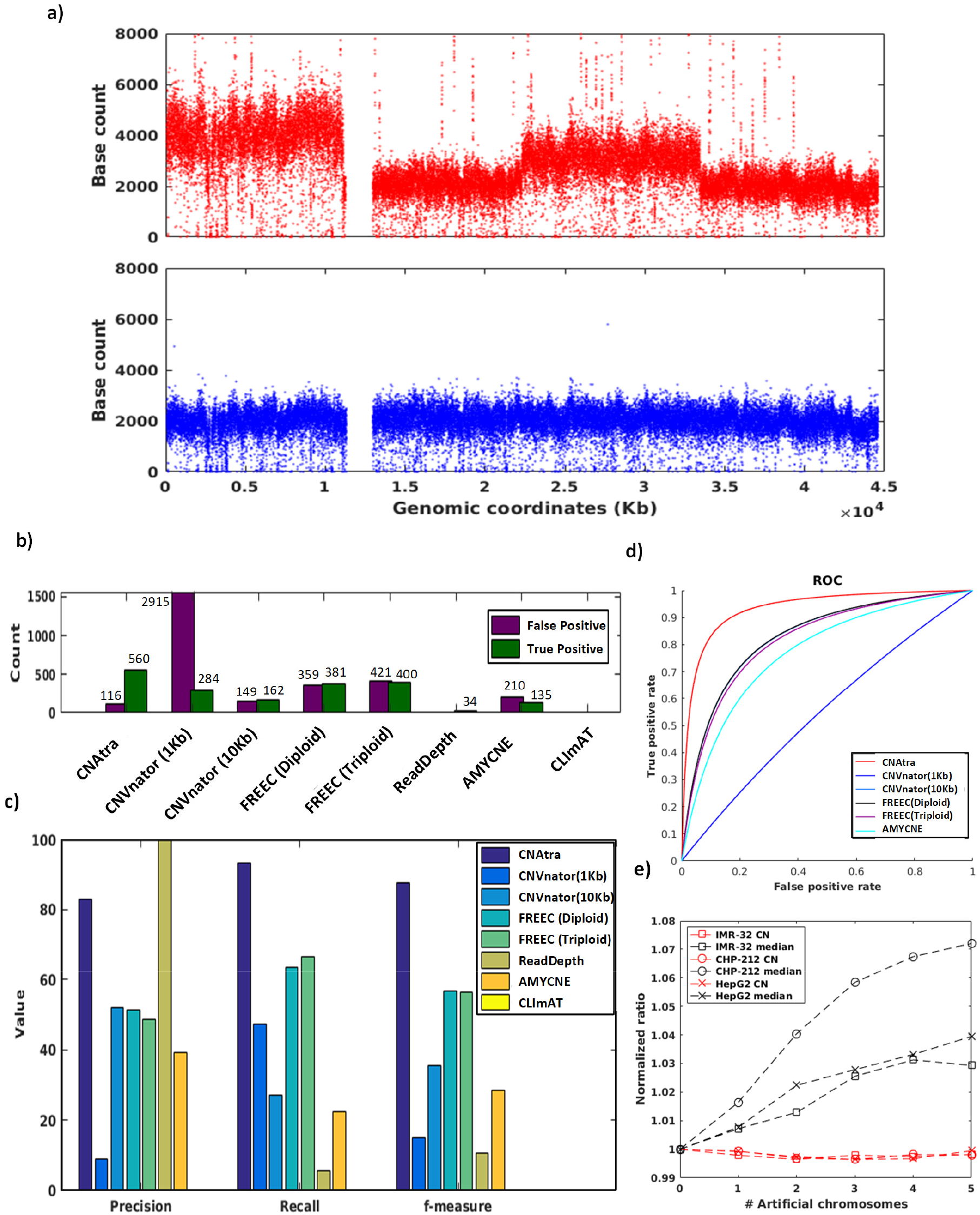
Performance evaluation of CNV detection tools on the simulated datasets. **(a)** Coverage plot of simulated data containing spiked-in LCVs and FAs (top panel). Coverage plot of CHP-212 Chr 12 (bottom panel) from which simulated data has been derived. **(b)** Bar graph showing false positive and true positive FAs detected by different tools. **(c)** Performance measurements (precision, recall and *f-measure*) of CNV detection tools for FAs. **(d)** Detection performances evaluation by the receiver operating characteristic (ROC) curves for FAs. **(e)** Line graph showing changes in CNAtra CN reference (red line) and the global median (black line) with each successive addition of artificial (simulated) chromosome in IMR-32 (square), CHP-212 (circle) and HepG2 (cross) cells.

We generated simulated CNV profiles using low-coverage (<2x) CHP-212, IMR-32, and HepG2 data. We introduced LCVs and focal CNVs in 5 chromosomes per cell line. For each chromosome, we have incorporated an average of 1-4 LCVs and then introduced 40 focal amplifications and deletions (size ranging between 50 and 100 kb) within these LCVs resulting in 600 FAs encompassing three cell lines. Upon introducing the CNAs, the overall RD signal maintains the multimodal characteristics of the cancer cells (Supplementary Fig. S6a). Using this simulated data, we benchmarked the performance of CNAtra against five RD-based single-sample CNV detection tools which include ReadDepth [14], CNVnator [9], FREEC [10], CLImAT [16] and AMYCNE [11]. We analyzed the performance of these tools after optimizing their parameters for low-coverage data (see Extended Methods under Supplementary Information). We set the criteria of >75% overlap between the spiked focal CNA and the CNV calls to be considered as a true call, and similarly, we set 90% overlap as criteria for true calls in case of LCVs. For a fair evaluation, we use the false amplifications only as false positives since many deletions may be falsely-identified at low-mappability regions and they affected by the mappability thresholding method which varies between different tools. Our evaluation showed that CNAtra could overall detect 93.3% (560/600) of the ‘introduced’ focal CNAs (IMR32 91.5%; CHP212 97% and HepG2 91.5%) (Fig. 4b; Supplementary Table 4). In comparison, the second best tool, FREEC can detect focal CNA with 63.5% (381/600) accuracy under diploid assumption. All other tools (CNVnator, AMYCNE, ReadDepth, and CLImAT) can detect focal CNVs with 0-50% accuracy (0-284 focal CNAs/600) (Supplementary Table 4). We also found that CNAtra has the highest accuracy for estimating copy number correctly with average CN difference of 0.251N followed by CNVnator with 10Kb bins (0.3642N) and FREEC (0.4357N) (Supplementary Fig. S6b). To evaluate the detection power, we computed precision and recall (sensitivity) of each tool and found that CNAtra outperforms all other tools.(Fig. 4c) Although ReadDepth showed the highest precision with zero false positives among all the tools, it can only detect 34/600 (5.6%) focal CNAs as 33 deletions and one amplification with wrong estimated CN. Therefore, we compute *f-measure* to estimate detection accuracy which balances the precision and the recall values. CNAtra showed the highest *f-measure* value of 87.77% followed by FREEC which showed 56.8% (Fig. 4c). Next, we plot the ROC (receiver operating characteristic) curve for evaluating the performance of the tools under the assumption that the maximum cumulative focal CNA locus length is 10% of the genome (for estimation of true negatives). The ROC curve clearly shows that CNAtra is superior in detecting focal CNAs in terms of true-positive and false-positive rates (Fig. 4d). When we compare the performance to detect LCVs, CNAtra again emerges at the top. CNAtra could detect 31/32 LCVs (96.8%) while CLImAT can detect 18/32 LCVs (56.25%) (Supplementary Table 4). Rest of the tools failed to detect any LCV event. We repeated this procedure to generate three additional simulated CNV profiles using different widths, frequencies, and copy numbers. All the analysis showed similar relative performance between the tools as demonstrative using ROC curves (Supplementary Fig. S6c; Supplementary Table 4).

In addition, we found that CNAtra is robust in estimating the CN reference regardless of the presence of LCVs. We stated earlier that LCVs could adversely affect local median that in turn can affect the copy number estimation. For example, after spiking the RD signal with LCVs, the global median changes by 3-7% (Fig. 4e) which may lead to the wrong estimation of CN reference. Despite this, CNAtra can correctly estimate the CN reference (Fig. 4e). We also analysed the computation time of all CNV detection tools (Supplementary Table 5). We have compared only the processing time for CNV calling modules since diferent tools have different input formats. We found that for low-coverage datasets, CNAtra, ReadDepth and CLImAT takes the shortest time.

### 3.8 Visualization and manual inspection of CNV calling exhibit CNAtra is best equipped to capture the complexity of cancer genomes

Review of CNA calls necessitates post-processing procedures which include visual inspection and curation of the results. Visualization and manual review of CNA profile in terms of copy number, size and structure can help to fine tune the tool parameters as well as refinement and curation of the results for downstream applications. Therefore, CNAtra provides an interactive visualization platform for the user to inspect and authenticate its results manually. We utilized this visual inspection approach to comprehensively understand the advantages and limitations of all tools using cancer datasets.

We found that CNAtra is the only tool to comprehensively detect both LCVs and focal alterations (Fig. 5; Supplementary Fig. S7 and S8). Moreover, we found that most tools are affected by imperfect segmentation of the large segments. For example, all tools except CNAtra have wrongly attributed the monoallelic 1p deletion in CHP-212 neuroblastoma cells into several segments. Only CLImAT identified this 1p deletion as a single event; however, they fail to correctly determine the right boundary of the segment (Fig. 5). Additionally, the focal amplification (FA1) inside the monoallelic 1p segmental deletion, which harbors enhancer region based on ENCODE ChromHMM [52], is detected by CNAtra, CNVnator and AMYCNE (Fig. 5). Also, 1q segmental amplification (correctly detected by CNAtra and CLImAT only) harbors many focal deletions. This confirms that focal amplification can be a part of the monoallelic segmental deletion and similarly focal deletions are present inside segmental amplification. None of the currently available tools addressed the coexistence of large and focal CNAs. Therefore, they cannot distinguish between these phenomena and tend to favor either of them. In addition, we found that all the tools suffer from the false estimation of copy number based on the proportion of LCVs in the genome. This results in misidentification of 1p and 1q regions with the same copy number by ReadDepth (Fig. 5). This effect is more evident in the A427 triploid cell line (Fig. 1d) [35]. As illustrated in Supplementary Figure 7, IB2 with CN 3 is misclassified as normal region and IB3 with CN 2 is wrongly identified as a deletion event since the median of the RD signal is matching the 3N (black line Fig. 1d A427) and not the correct CN reference (2N) (blue line in Fig. 1d A427). All the tools are affected by overdispersion in low-coverage data which may result in false positives and false negatives as estimated using simulated data. CNAtra circumvents this problem using thresholding parameters; however, the user can apply higher stringency thresholding to curate the CNA data manually.

**Fig. 5.**
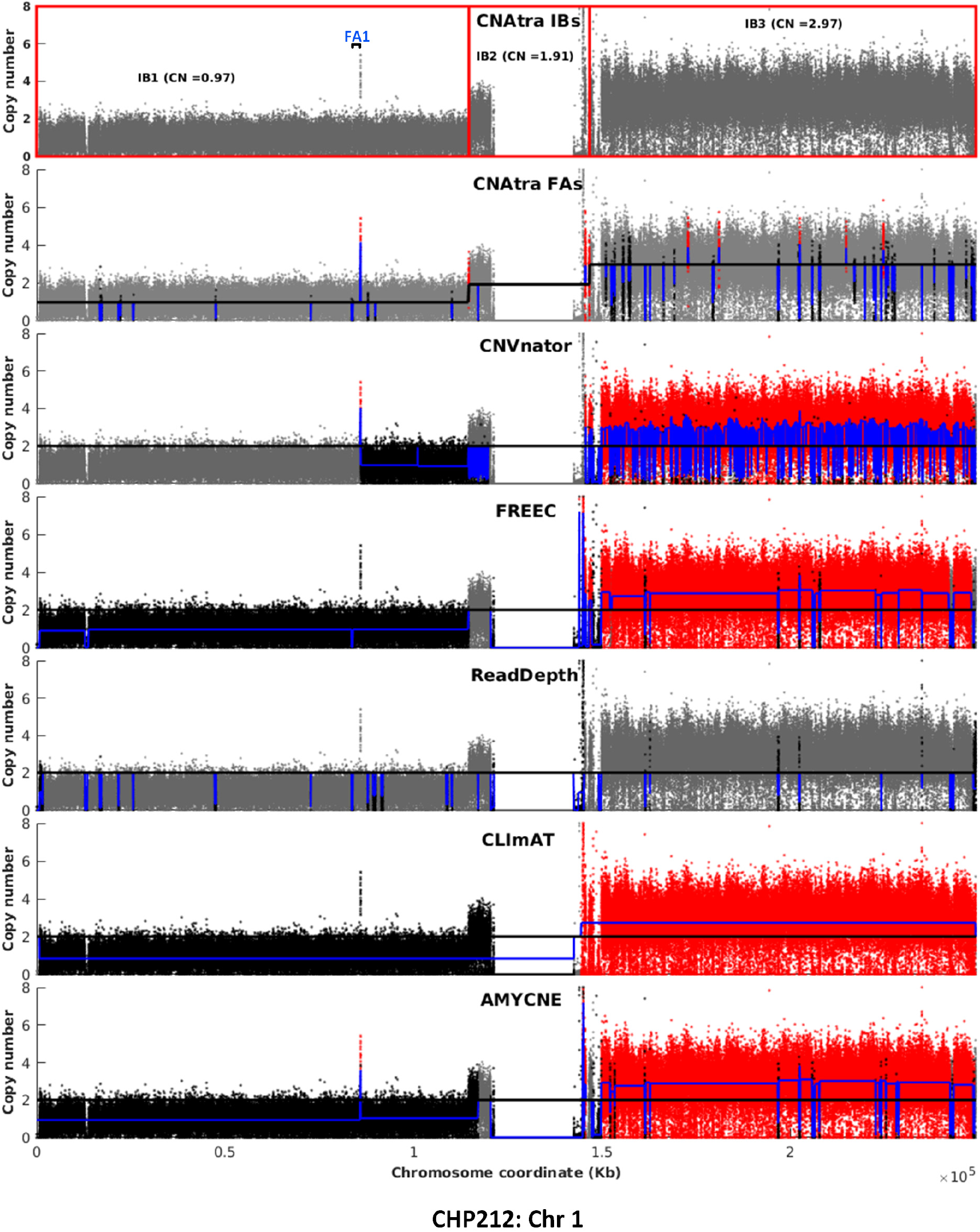
Visual comparison of CNA profiles generated by CNV detection tools on Chr 1 of CHP-212. Red, black, grey dots are bins belonging to focally amplified, deleted, and CN neutral regions respectively. The blue line represents the copy number of each CNA. Any amplitude transition indicates a new CNA region. Top panel shows the IBs identified by CNAtra (each red box represents one IB). CNAtra examines each IB to detect the focal CNVs (second panel). Rest of the panels show CNV output by other detection tools.

## 4. DISCUSSION

Hyperploidy and pervasive genetic alterations are the hallmark of cancer genomes. In the current study, we analyzed several cancer cells with different levels of aneuploidy which showed a complex multimodal distribution due to widespread large-scale and focal CNAs. This is in stark contrast to unimodal RD signal distribution of normal human genomes (1000 genomes project) devoid of large segmental aneuploidies. Current CNV detection tools have limited ability to handle cancer genomes due to their assumption of unimodal probabilistic distributions of RD signal, Erroneous modeling of the RD signal distribution may lead to incorrect estimation of CN reference and false segmentation which adversely affects the final CNA results. Consequently, cancer CNA profiles available from public databases (*viz*. COSMIC, CCLE) face the same problem as their calls are based on a similar assumption. CNAtra successfully utilized a multimodal distribution to estimate the CN reference (global reference) and then employed a CN-based merging algorithm first to detect the large segments. Then each CN-designated segment formed the basis for detecting focal alteration(s) where its copy number is used as a local reference.

High-coverage (>15x) WGS datasets are generally used for CNA profiling; however, they are not available for many cancer cell lines. As an alternative solution, the ChIP-seq ‘input’ NGS reads can be effectively used which are publicly available for many cell lines. These input reads are generated from sonicated chromatin and typically used to normalize and peak calling of the ChIP-seq data. These input data contain the genome-wide reads but they are generally sequenced at low-coverage (<2x). These data can be judiciously utilized to compute the RD signal for CNA analysis. Any low-coverage dataset is afflicted with overdispersion of the RD signal which prompts detection of a high number of false positives. CNAtra bypasses this problem by using coverage-based thresholding to detect highly significant focal alterations minimizing false positives. We have estimated the relation between coverage and thresholds using subsampling of four high-coverage data including one cancer data. Our heuristic approach of determining coverage-based thresholding parameters leaves space for future improvements by incorporating more high-coverage datasets. For example, the dispersion levels vary based on the CN state of each LCVs. Therefore, training using more datasets will provide better estimation of thresholding parameters at different CN states. Also, presence of poor-mappability regions (bins) can lead to false detection of homozygous focal deletions surrounding these bins. CNAtra provides the option to filter these focal deletions based on the percentage of low-mappability bins.

One of the major limitations of the performance evaluation of CNV detection tools is the non-availability of a complete repertoire of ‘experimentally validated’ cancer CNA profiles. Therefore, we generated realistic simulated data using the available cancer data as input to maintain the signal variability and features of cancer RD signal. Then we introduced CNAs of different length-scales randomly as ‘artificial’ ground truth. We benchmarked CNAtra against five currently available tools using this simulated data. We established that CNAtra is the only tool to stratify large-scale and focal CNAs which reflect critical biological features. CNAtra beats all other competing tools in higher precision and sensitivity which are corroborated by visual inspection. To sum up, we believe that CNAtra is the ideal approach (guide) to tackle complex and low-coverage cancer datasets. CNAtra has immense potential to add value towards the study of cancer genomes as well as discovery of novel CNAs.

## Declarations

### Acknowledgment

We acknowledge Sanyal and Chattopadhyay lab members for their valuable comments.

### Author’s contributions

AS, AC and AISK conceived the project. AISK developed CNAtra software with inputs from AS and AC, performed all the analyses. CK helped in the read filtering, RD signal calculation as well as installing and running available CNV detection tools. AISK and CK developed the simulated data with inputs from AS. AS, AISK and AC analyzed the data and prepared the manuscript. All authors read and approved the final manuscript.

### Ethics approval and consent to participate

Not applicable.

### Supplementary data

Supplementary Information containing Extended Methods and Supplementary Figures are provided as Supplementary data. Supplementary Tables and CNV results of all CNV detection tools are provided under Additional files.

